# Unsupervised cluster analysis of SARS-CoV-2 genomes reflects its geographic progression and identifies distinct genetic subgroups of SARS-CoV-2 virus

**DOI:** 10.1101/2020.05.05.079061

**Authors:** Georg Hahn, Sanghun Lee, Scott T. Weiss, Christoph Lange

## Abstract

Over 10,000 viral genome sequences of the SARS-CoV-2 virus have been made readily available during the ongoing coronavirus pandemic since the initial genome sequence of the virus was released on the open access Virological website (http://virological.org/) early on January 11. We utilize the published data on the single stranded RNAs of 11, 132 SARS-CoV-2 patients in the GISAID (Elbe and Buckland-Merrett, 2017; Shu and McCauley, 2017) database, which contains fully or partially sequenced SARS-CoV-2 samples from laboratories around the world. Among many important research questions which are currently being investigated, one aspect pertains to the genetic characterization/classification of the virus. We analyze data on the nucleotide sequencing of the virus and geographic information of a subset of 7, 640 SARS-CoV-2 patients without missing entries that are available in the GISAID database. Instead of modelling the mutation rate, applying phylogenetic tree approaches, etc., we here utilize a model-free clustering approach that compares the viruses at a genome-wide level. We apply principal component analysis to a similarity matrix that compares all pairs of these SARS-CoV-2 nucleotide sequences at all loci simultaneously, using the Jaccard index (Jaccard, 1901; Tan et al., 2005; Prokopenko et al., 2016; Schlauch et al., 2017). Our analysis results of the SARS-CoV-2 genome data illustrates the geographic and chronological progression of the virus, starting from the first cases that were observed in China to the current wave of cases in Europe and North America. This is in line with a phylogenetic analysis which we use to contrast our results. We also observe that, based on their sequence data, the SARS-CoV-2 viruses cluster in distinct genetic subgroups. It is the subject of ongoing research to examine whether the genetic subgroup could be related to diseases outcome and its potential implications for vaccine development.

## 1 Introduction

Now, the outbreak of severe acute respiratory syndrome coronavirus 2 (SARS-CoV-2) has created a pandemic as it has spread swiftly from continent to continent, resulting in >2,300,000 infections, with approximately 5% mortality and a devastating effect on public health. Little is known about the rates of mutation in SARS-CoV-2, viral genetic variation is generated on the same timescale as virus transmission which allows us to track the spread of SARS-CoV-2. Phylogenetic and phylogeographic analysis conferred insights into distinct subtypes as the virus spreads through the global population and could be helpful in untangling complex virus transmission dynamics and understanding disease susceptibility (Mousavizadeh and Ghasemi, 2020).

One of the striking features of SARS-CoV-2 infection is the variable clinical features of the disease. In general, older persons seem to have greater severity and mortality (Zhou et al., 2020), as do males (Shi et al., 2020). However, some younger subjects have had fatal disease as well (Zhou et al., 2020). Organ involvement also varies with respiratory complications being most frequent but cardiac and renal complications occurring in some other patients (Zhou et al., 2020). This degree of clinical variation may be due to environmental factors such as confinement to a nursing home or chronic care facility (McMichael et al., 2020), or genetics of either the host or the virus. While studies of host genetics are underway, we have chosen here to focus on genetic variation of the virus.

By mid June 2020, the GISAID (Elbe and Buckland-Merrett, 2017; Shu and McCauley, 2017) database contained samples of the sequenced SARS-CoV-2 genome obtained from thousands of individuals infected with the human coronavirus disease. After routine cleaning and alignment of the data (see the *Methods* section for details), we were left with *n* = 7, 640 samples having a trimmed length of *p* = 28, 784 nucleotide sequences. In the principal component cluster analysis, we focused on the loci in the SARS-CoV-2 genome where differences to the SARS-CoV-2 reference sequence (in Hamming distance) are observed. The reference sequence is available on GISAID under the accession number EPI ISL 412026. The Hamming matrix indicates the mismatches to the reference sequence, and its row sums are the Hamming distance of each nucleotide sequence to the reference sequence.

As we are interested in the geographical/temporal change of the SARS-CoV-2 genome, we assessed the similarity between each pair of virus genomes with the Jaccard index (Jaccard, 1901; Tan et al., 2005). By computing the Jaccard index for each pair of samples based on all loci we constructed the Jaccard similarity matrix for the 7, 640 virus genomes (Prokopenko et al., 2016; Schlauch et al., 2017). The Jaccard similarity matrix can efficiently be computed across a large set of genomes and is a powerful tool to detect genetic substructures/clusters in such sets (Hahn et al., 2020c,b).

The article is structured as follows. Section 2 describes the preparation of the dataset including cleaning and alignment, as well as the Jaccard similarity matrices we compute. Clustering results for the SARS-CoV-2 genome samples are presented in Section 3, where they are contrasted with a phylogenetic analysis. The article concludes with a discussion in Section 4.

## 2 Methods

### 2.1 Data acquisition

All analyses are based on nucleotide sequence data downloaded from GISAID (Elbe and Buckland-Merrett, 2017; Shu and McCauley, 2017) on 14 June 2020. Only complete sequences (defined as having a length ≥ 29000) were selected, resulting in 45, 895 sequences.

### 2.2 Data curation

We removed sequences with reading errors in the *fasta* file (defined as letters other than A, C, G, T). Carrying out an exhaustive search for a reference sequence found a sequence of length 300 which we used as an anchor to align all samples. For this, we exhaustively enumerated all subsequences of the nucleotide sequences in our sample, and checked for each one of them if it is found at roughly the same position in all samples. The size of 300 bp was chosen arbitrarily; however, this choice allowed us to accurately align all samples.

AATGTATACATTAAAAATGCAGACATTGTGGAAGAAGCTAAAAAGGTAAA ACCAACAGTGGTTGTTAATGCAGCCAATGTTTACCTTAAACATGGAGGAG GTGTTGCAGGAGCCTTAAATAAGGCTACTAACAATGCCATGCAAGTTGAA TCTGATGATTACATAGCTACTAATGGACCACTTAAAGTGGGTGGTAGTTG TGTTTTAAGCGGACACAATCTTGCTAAACACTGTCTTCATGTTGTCGGCC CAAATGTTAACAAAGGTGAAGACATTCAACTTCTTAAGAGTGCTTATGAA

We grouped samples together depending on the location tag in their *fasta* file using the following nine categories: (1) China (all provinces apart from Wuhan), (2) Wuhan, (3) Korea, (4) Europe (all countries apart from Italy), (5) Italy, (6) Africa, (7) Australia and New Zealand, (8) South(east) Asia, (9) North America (USA, Canada), (10) Central America (including Mexico), (11) South America, (12) Japan, and (13) Iran. Selecting only samples falling into those regions, having a complete time stamp (year, month, date), and having an exact match of the above reference sequence resulted in *n* = 7, 640 samples.

### 2.3 Trimming and comparison to reference sequence

After alignment of all *n* samples with the anchor, we trimmed sequences to the left and right in order to establish a sequence window in which all sequences had reads. This window had a sequence length of *p* = 28, 784.

We denote with *X* ∈ 𝕊^*n*×*p*^ the matrix of nucleotide sequences. Letting *r* ∈ 𝕊^*p*^ be the reference sequence published on GISAID (accession number EPI ISL 412026), where 𝕊 = {*A, C, G, T*}, we compute a Hamming matrix *Y* ∈ {0, 1}^*n*×*p*^ by defining *Y*_*ij*_ = 1 if and only if *X*_*ij*_ ≠*r*_*j*_, and *Y*_*ij*_ =0. otherwise. The matrix *Y* indicates the mismatches to the reference sequence, and its row sums are the Hamming distance of each nucleotide sequence to the reference sequence.

### 2.4 Measuring similarity

We employ the locStra (Hahn et al., 2020c,b) R-package to calculate the Jaccard similarity matrix of *Y*, denoted as *J* :=*Jac*(*Y*) ∈ ℝ^*n*×*n*^. The Jaccard matrix measures the similarity between all pairs of the *n* curated samples from GISAID. Computing the first two eigenvectors of *J* (for plotting) completes the numerical part of the analysis.

## 3 Results

Figure 1 displays the first two principal components of the Jaccard matrix for the analyzed SARS-CoV-2 genomes, labelled with the country information obtained from GISAID. We observe four distinct branches/clusters for both European and North American samples (Fig.1, left). The top branch contains most cases from the US, the branch below is a mixture of US and European cases. The horizontal branch contains predominantly European cases, while the bottom branch contains a mixture of samples from Europe, China, and South(east) Asia. The predominantly European branch on the x-axis seems to be linked to the cluster at the origin, by genomes from Wuhan, Italy and Europe (Fig. 1, right).

**Figure 1:**
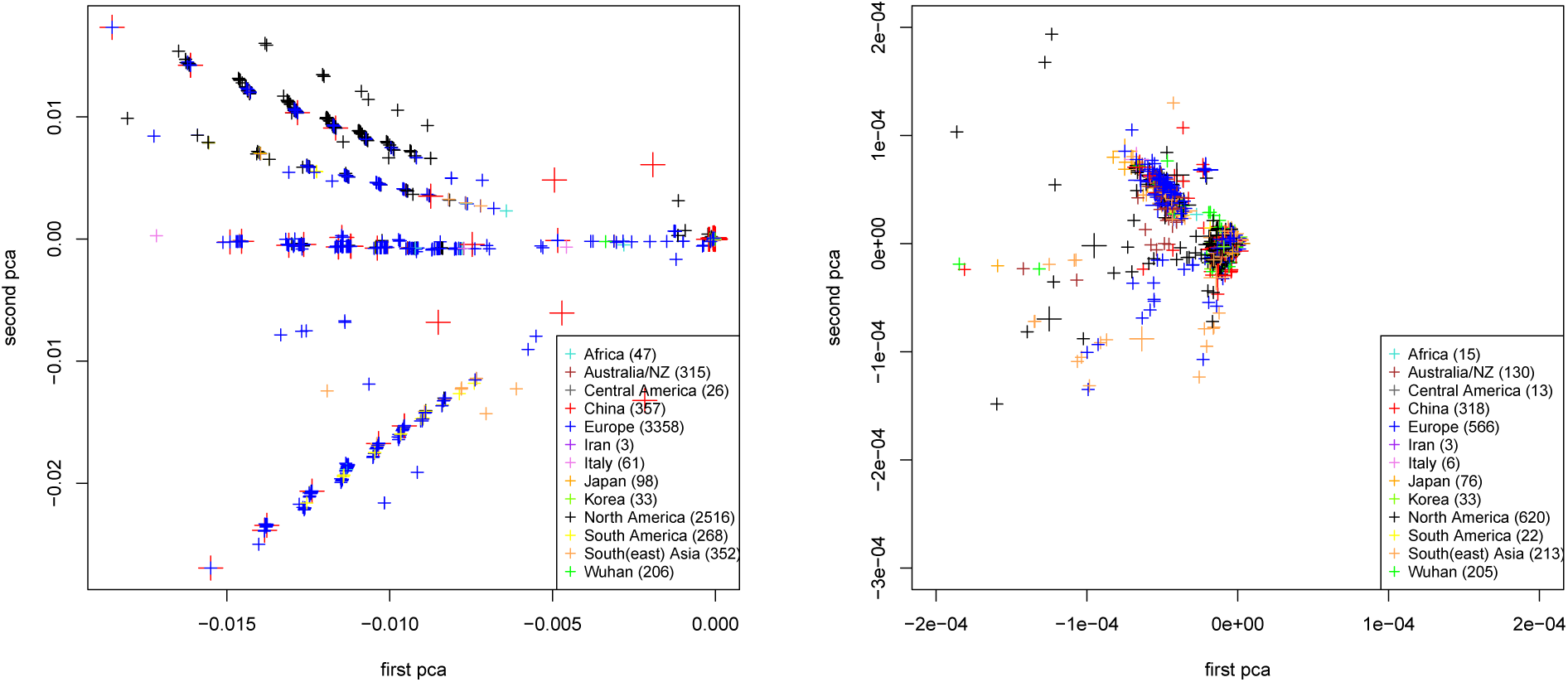
First two principal components of the Jaccard similarity matrix for the 7, 640 SARS-CoV-2 genomes by region/country. Entire dataset (left) and zoomed-in region around the origin (0, 0) (right). Numbers in brackets for each country denote the number of SARS-CoV-2 genomes which are visible in each plot.

Furthermore, we observe a dense cloud of points around the origin (0, 0), which contains indistinguishable data points for genomes from all countries of the data set. When looking at the zoomed-in region in Fig. 1 (right), we observe that starting from the origin (0, 0), which has the highest density of genomes from China and Wuhan, a spatial spread-out is seen in which genomes from geographically close locations (Korea and South/Southeast Asia) cluster in the neighborhood of the origin. This “original” cluster contains also a relatively large number of virus genomes from the US and Europe, which could suggest early transmissions to these geographic regions. Almost all of the data points for the genomes from Wuhan and China are part of this cluster around the origin (0, 0), reflecting where SARS-CoV-2 was first reported.

For a better understanding of the distinct clustering of the European and US samples into the four observed subbranches (Fig. 1, left), we examine the genomic locations where the four subgroups differ. In particular, we are interested in how SARS-CoV-2 samples belonging to those four branches compare to the SARS-CoV-2 reference sequence. For the reference genome subdivided into 50 consecutive bins, Fig. 2 shows the normalized numbers of mismatches when comparing the trimmed reference sequence to the four branches of European and American samples. The number of mismatches is normalized with respect to the bin size and the number of samples in each branch. We observe that, besides of the (outlier) mismatches at the beginning and at the end of the virus genome, the samples of the four distinct branches differ from the reference genome at the same four genomic regions, and by roughly similar frequencies. The only exception is the fourth branch at approximately position 25000, where the first two branches are similar, and the third and fourth branches are similar.

**Figure 2:**
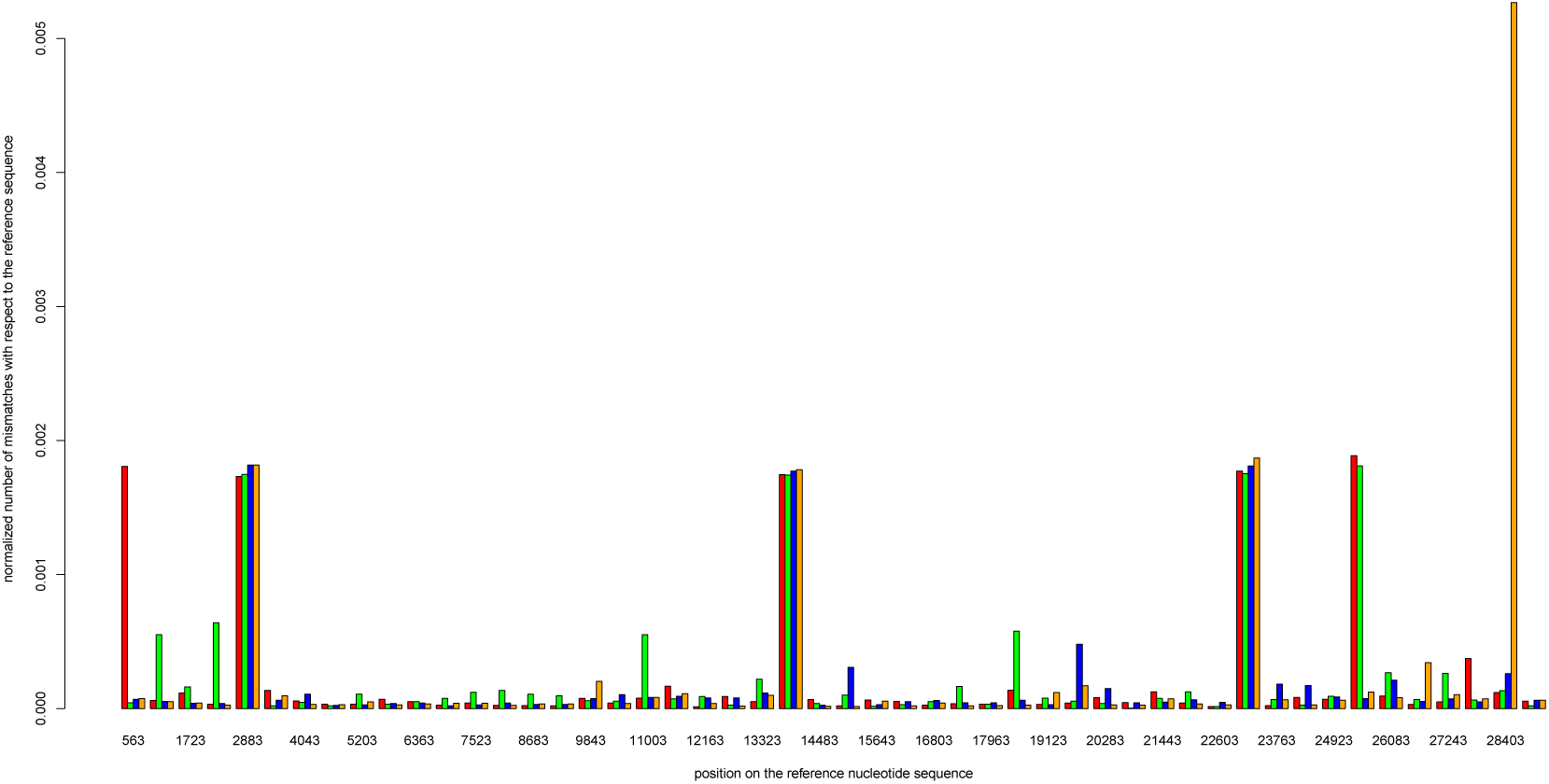
The x-axis shows the roughly 29, 000 nucleotides of the trimmed SARS-CoV-2 reference sequence in 50 bins. The y-axis shows per bin the normalized number of mismatches (with respect to the reference sequence) among the samples in the European and North American population, stratified into samples from the top branch (red), the second branch (green), the middle branch (blue), and the bottom branch (orange) visible in the left panel of Fig. 1. The normalization is done with respect to both the bin size and the number of samples in each branch.

We also aim to contrast our results with the ones of a phylogenetic analysis, computed with the “TreeTool App” on the GISAID website (Freunde of GISAID, 2020) for GISAID samples falling into the same time range as the ones used in our analysis. The analysis with the TreeTool App is based on an alignment with MAFFT (Katoh, 2013), from which a phylogenetic tree is subsequently built using the tool “FastTree” (Price et al., 2010; Price, 2020). A comparison of our clustering in Figure 1 with the phylogenetic analysis in Figures 3 and 4 reveals notable similarities and differences.

**Figure 3:**
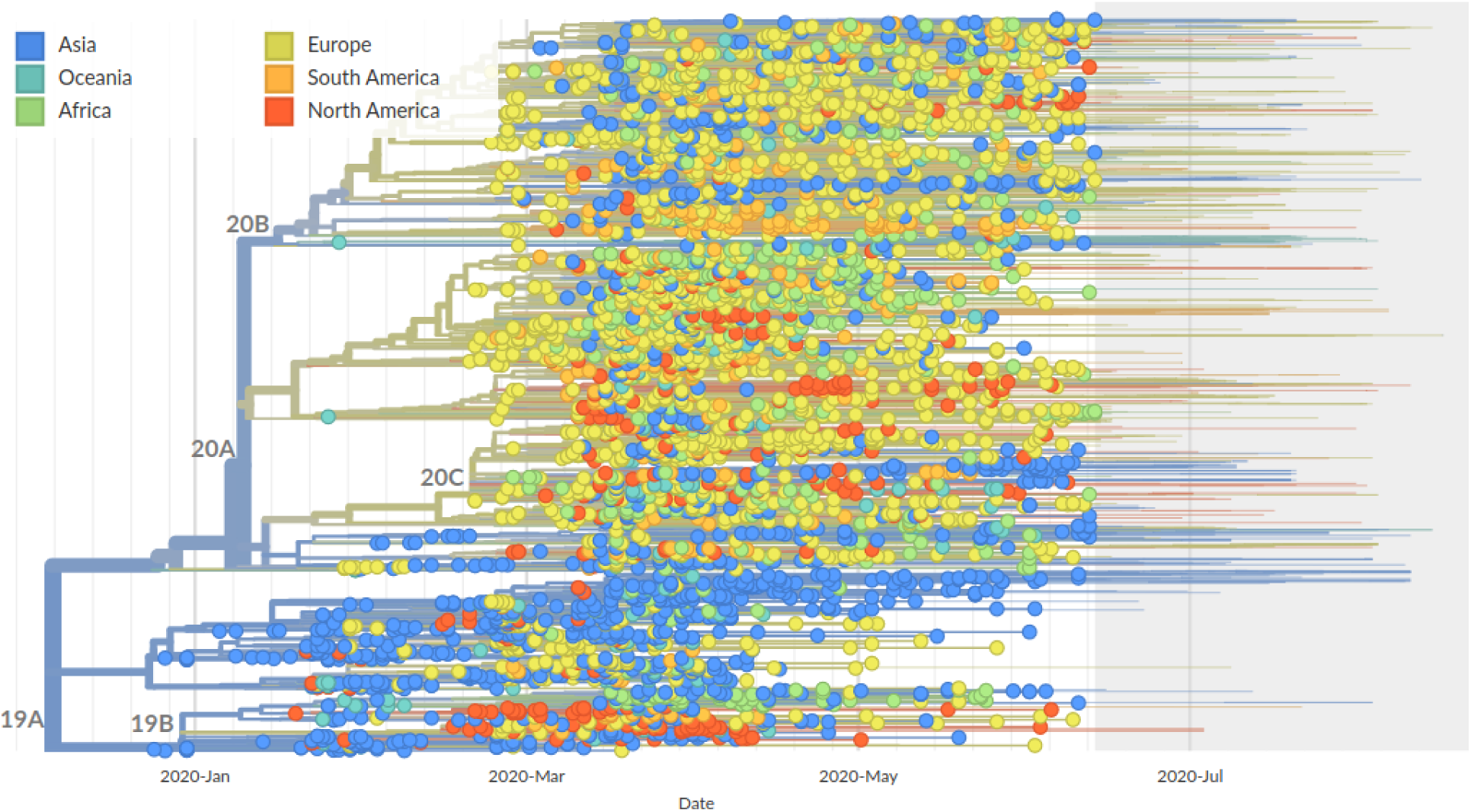
Phylogenic analysis as tree computed on https://www.gisaid.org/ epiflu-applications/influenza-phylogenetics/ using MAFFT for sequence alignment and FastTree for building the phylogenic tree.

**Figure 4:**
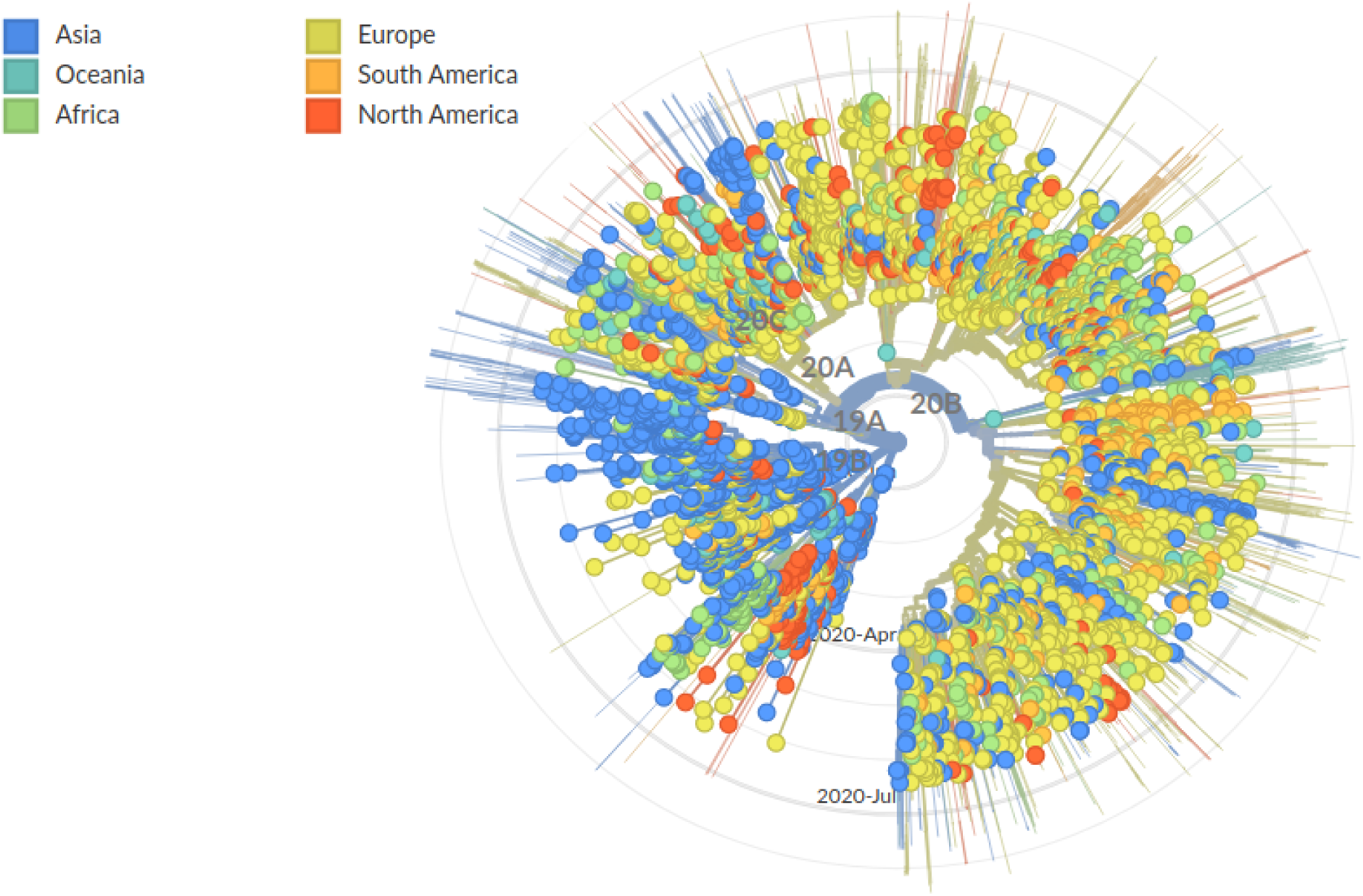
Phylogenic analysis as radial representation computed on https://www.gisaid.org/ epiflu-applications/influenza-phylogenetics/ using MAFFT for sequence alignment and FastTree for building the phylogenic tree.

We observe that, likewise to our approach, Figure 3 reveals four clusters (which emerge in roughly March 2020). The bottom one contains mostly a mix of Asian samples, similarly to the bottom cluster we observe. The second cluster (from the bottom) contains a mix of Asian and North American/ European samples, the third is mostly a mix of North American and European samples, and the top cluster contains mostly Asian and European samples. Here, the clusters in the center of the plot correspond to the middle cluster of mostly European samples that we observe in Figure 1. However, in contrast to Figure 3, our clustering analysis shows a stronger mixing of mostly European and North American samples in the top two clusters in Figure 1, whereas such a result is not observed for the phylogenetic tree.

While both approaches, principal components and phylogenetics, identify the same global clustering structures, the differences in the analysis results between the approaches are observed in the local clustering substructure. Given the limited data, the question which of the two approaches is preferable cannot be answered conclusively at this point. Amongst others, this decision will depend on the specifics of the application, the analysis question, and further verification of the results from either approach through external findings.

## 4 Discussion

Our model-free, global analysis of the SARS-CoV-2 sequences demonstrates that the genome of the virus varies by geographic region, with different viral sequences present in Asia vs. Europe and the US. Our analysis is complementary to model-based approaches, e.g. phylogenetic tree approaches, or estimation of mutation rates (Zhao et al., 2004). Our analysis results also suggest that there are four distinct genetic subgroups in Europe and the US. The viral sequences that were initially observed in Asia seem to be more homogeneous. The genetic sequences of viruses from Europe and the US are more diverse when compared to the sequences from countries in Asia. If new cases are observed, e.g. new cases in Beijing in June 2020 (Xiaohua, 2020; Liu and Nebehay, 2020), clustering approaches can help to understand the origin of the transmissions, as one can examine whether the genomes of the new cases belong to a previously observed cluster (Hahn et al., 2020a).

It is important to consider the methodological aspects of our analysis to describe the viral genetic diversity of SARS-CoV-2. We used a direct, unsupervised approach to compare the entire viral genomes, i.e. principle components analysis based on Jaccard similarity matrices. We did not include information about the geographic origin of the samples in the analysis nor did we attempt to directly model the evolutionary relationship of the different SARS-CoV-2 genomes, e.g. via phylogenetic analysis. Nevertheless, our analysis results reflect clearly the chronological spread of SARS-CoV-2 around the globe. Furthermore, the principal components that are obtained by our approach can be tested for dependency/association with other important variables, e.g. time. Moreover, while our approach currently does not allow for the incorporation of genomes with missing information, is our ongoing methodological work to allow for such an analysis. Both aforementioned directions are work in progress.

The advantages of our methodology compared to existing analyses are twofold: First, our unsupervised approach is capable of recovering geographic subgroups and geographic progression without any (spacial) information, and it takes all loci into account simultaneously. Second, it is particularly simple as it only involves the Hamming matrix and its Jaccard representation, both of which can be efficiently computed using binary operations only (Hahn et al., 2020b).

To be precise, the computational effort existing software such as “FastTree” (Price, 2020) is of the order of *O*(*n*^1.5^ log(*n*)*p*) (Price et al., 2010), where *n* is the number of sequences and *p* is the length of the alignment. In our approach, the computation of the Hamming matrix can be done in *O*(*np*), since it requires only the entry-wise comparison of the aligned reference genome to each trimmed nucleotide sequence. Afterwards, the computation of the Jaccard matrix on the Hamming matrix requires effort *O*(*spn*^2^), see Hahn et al. (2020b), where *s* is the sparsity of the input (Hamming) matrix. For the case of the SARS-CoV-2 analysis presented in this work, it turns out that after alignment, all SARS-CoV-2 sample sequences naturally share a lot of nucleotides, which is reflected in a proportion of non-zero entries of only 0.0067 for the GISAID data we analyze. The high sparsity effectively brings down the effort to an empirical effort of *O*(*n*^1.5^*p*), thus making our approach indeed more efficient than FastTree. For approximate computations, the calculation of the Jaccard matrix can be avoided altogether, leaving only the computation of the Hamming matrix in *O*(*np*), and an *O*(*np*) effort to calculate the first two eigenvectors using the technique of (Halko et al., 2011, Section 1.6).

Methodology to identify clusters in genomic data, using combination of discriminant analysis and prior dimensionality reduction via PCA, could be applied as future work (Jombart et al., 2010). However, the main goal of our submission is to show that a simple approach based on Hamming distance and Jaccard similarity matrices reveals a meaningful stratification of SARS-CoV-2 genomes. Other approaches such as Lemey et al. (2020) focus on the incorporation of travel history data into the (phylogenetic) analysis, which is demonstrated for a small number of genomes. Such an approach can be more informative, however can be difficult to obtain travel history for thousands of samples. Here, we aimed instead to reflect a geographic progression and spatial clustering of SARS-CoV-2 based solely on genomic data.

Our analysis results have two potentially important implications. It is the subject of ongoing research to understand whether the four genetic subgroups of the SARS-CoV-2 viruses have different clinical implications in terms of disease progression and disease outcome.

One important research question in this context is how to link the sequenced viral genome information to clinical outcomes of Covid-19. In this context, we believe that the PCA clustering methodology could provide new insights as it enables the application of all the available methodologies that have been developed for genome-wide association studies (GWAS) in human genetics. For instance, one could conduct a genome-wide association study for Covid-19 in which the loci of the viral genomes are tested for association with (that is, regressed on) the clinical outcome variable of the host/patient. To guard such an analysis against confounding/substructure in the viral genome data, one can include the eigenvectors of the PCA analysis into the GWAS regression models. Initial applications of the GWAS methodology to sequenced SARS-CoV-2 genomes and their host’s/patient’s clinical data have identified loci in the SARS-CoV-2 genome that could be associated with mortality (Lange et al., 2020). Phylogenetic approaches do not necessarily offer this convenience.

Furthermore, our results suggest that vaccine creation will have to take into account the genetic variability in viral sequence described here. As the virus has spread over time, its genetic diversity has increased and is likely to increase even further. To combat the virus successfully, it will be fundamental to understand the consequences of this development on the features of the virus.

## Acknowledgements

The authors gratefully acknowledge the contributors, originating and submitting laboratories of the sequences from GISAID’s EpiCoV™ Database (Elbe and Buckland-Merrett, 2017; Shu and

McCauley, 2017) on which this research is based. A detailed list of contributors is available in the Supplementary Information.

## Data Availability Statement

Sequence data that support the findings of this study are deposited in the GISAID database with accession numbers in the range of EPI ISL 402119 to EPI ISL 467430 (https://www.gisaid.org/).

